# Whole transcriptome profiling of placental pathobiology in SARS-CoV-2 pregnancies identifies placental dysfunction signatures

**DOI:** 10.1101/2023.01.20.524893

**Authors:** Nataly Stylianou, Ismail Sebina, Nicholas Matigian, James Monkman, Hadeel Doehler, Joan Röhl, Mark Allenby, Andy Nam, Liuliu Pan, Anja Rockstroh, Habib Sadeghirad, Kimberly Chung, Thais Sobanski, Ken O’Byrne, Florido Almeida Ana Clara Simoes, Patricia Zadorosnei Rebutini, Cleber Machado-Souza, Emanuele Therezinha Schueda Stonoga, Majid E Warkiani, Carlos Salomon, Kirsty Short, Lana McClements, Lucia de Noronha, Ruby Huang, Gabrielle T. Belz, Fernando Souza-Fonseca-Guimaraes, Vicki Clifton, Arutha Kulasinghe

## Abstract

**Objectives:** Severe Acute Respiratory Syndrome Coronavirus 2 (SARS-CoV-2) virus infection in pregnancy is associated with higher incidence of placental dysfunction, referred to by a few studies as a “preeclampsia-like syndrome”. However, the mechanisms underpinning SARS-CoV-2-induced placental malfunction are still unclear. Here, we investigated whether the transcriptional architecture of the placenta is altered in response to SARS-CoV-2 infection.

**Methods:** We utilized whole-transcriptome, digital spatial profiling, to examine gene expression patterns in placental tissues from participants who contracted SARS-CoV-2 in the third trimester of their pregnancy (n=7) and those collected prior to the start of the coronavirus disease 2019 (COVID-19) pandemic (n=9).

**Results:** Through comprehensive spatial transcriptomic analyses of the trophoblast and villous core stromal cell subpopulations in the placenta, we identified signatures associated with hypoxia and placental dysfunction during SARS-CoV-2 infection in pregnancy. Notably, genes associated with vasodilation (*NOS3*), oxidative stress (*GDF15*, *CRH*), and preeclampsia (*FLT1, EGFR, KISS1, PAPPA2),* were enriched with SARS-CoV-2. Pathways related to increased nutrient uptake, vascular tension, hypertension, and inflammation, were also enriched in SARS-CoV-2 samples compared to uninfected controls.

**Conclusions:** Our findings demonstrate the utility of spatially resolved transcriptomic analysis in defining the underlying pathogenic mechanisms of SARS-CoV-2 in pregnancy, particularly its role in placental dysfunction. Furthermore, this study highlights the significance of digital spatial profiling in mapping the intricate crosstalk between trophoblasts and villous core stromal cells, thus shedding light on pathways associated with placental dysfunction in pregnancies with SARS-CoV-2 infection.

**Graphical abstract:** In this study, using spatial digital profiling transcriptomic approaches, we demonstrate that SARS-CoV-2 infection in pregnancy disrupts optimal placental function by altering the genomic architecture of trophoblasts and villous core stromal cells.

## INTRODUCTION

Viral infections in pregnancy can disrupt placental function and predispose pregnancy complications, including late-onset preeclampsia, preterm birth, stillbirth, and intrauterine fetal demise^1–4^. Recent studies have revealed that pregnant women who contract Severe Acute Respiratory Syndrome Coronavirus 2 (SARS-CoV-2, which causes coronavirus disease 2019 [COVID-19]), can experience placental dysfunction and what has been referred to as “preeclampsia-like syndrome” ^5–10^. Placental tissues from COVID-19 patients exhibit increased vasculopathy and inflammation, which are characteristic pathological features of preeclampsia ^11^. Moreover, clinical manifestations observed in COVID-19 patients, such as COVID-19-associated hypoxia, hypertension, endothelial dysfunction, kidney disease, thrombocytopenia, and liver injury, overlap with those observed in preeclampsia^5, 12^. However, mechanisms through which SARS-CoV-2 infection predisposes pregnancies to these preeclampsia-like pathological features are largely unclear.

The placenta is vital for fetal development and growth throughout gestation as it is a functional interface between the mother and fetus^13^. This interface comprises various anatomically distinct sites, including the decidua basalis, where maternal immune cells and decidual stromal cells interact with fetal extravillous trophoblasts^14^. The maternal-fetal interface also consists of the placental intervillous space, where maternal immune cells interact with fetal syncytiotrophoblasts, and the boundary between the parietalis and the chorion laeve in the chorioamniotic membranes^14^. Other cell types within this interface, such as villous cytotrophoblasts, column cytotrophoblasts, fibroblasts, endothelial cells, and Hofbauer cells, contribute to nutrient and waste exchange, hormone production, protection from pathogens, and maternal immune responses essential for fetal development ^13, 14^. Whether SARS-CoV-2 infection modifies the transcriptomic architecture and functional characteristics of different cell types within these distinct placental sites is still unclear.

In this study, we utilised digital spatial whole-transcriptomic analysis of human placental tissue to delineate molecular pathways associated with SARS-CoV-2 infection-induced placental pathology in pregnancy. Specifically, we focused our analysis on defining the distinct transcriptional profiles of trophoblasts and villous core stromal cell populations (the latter including endothelial, fibroblast, and immune cells), in the context of SARS-CoV-2 infection. We identified several pivotal pathways that underlie the development of a “preeclampsia-like syndrome” associated with SARS-CoV-2 infection in pregnancy.

## RESULTS

### Characterization of patient demographics and histopathology in collected placentae

Tissue microarrays were constructed using placental cores that were collected immediately after birth from unvaccinated participants who had tested positive within 15 days prior to delivery (Alpha strain, April 2020, n=7), and unvaccinated participants who were negative for SARS-CoV-2 throughout their pregnancy (n=9; Table 1). There were no significant differences in placental weight, fetal weight, gestational age, comorbidities, or maternal age between the two groups (Table 1). Within the SARS-CoV-2 group, 3/7 newborns were born preterm, compared to 4/9 in the control group (Table 1).

**Table 1:**
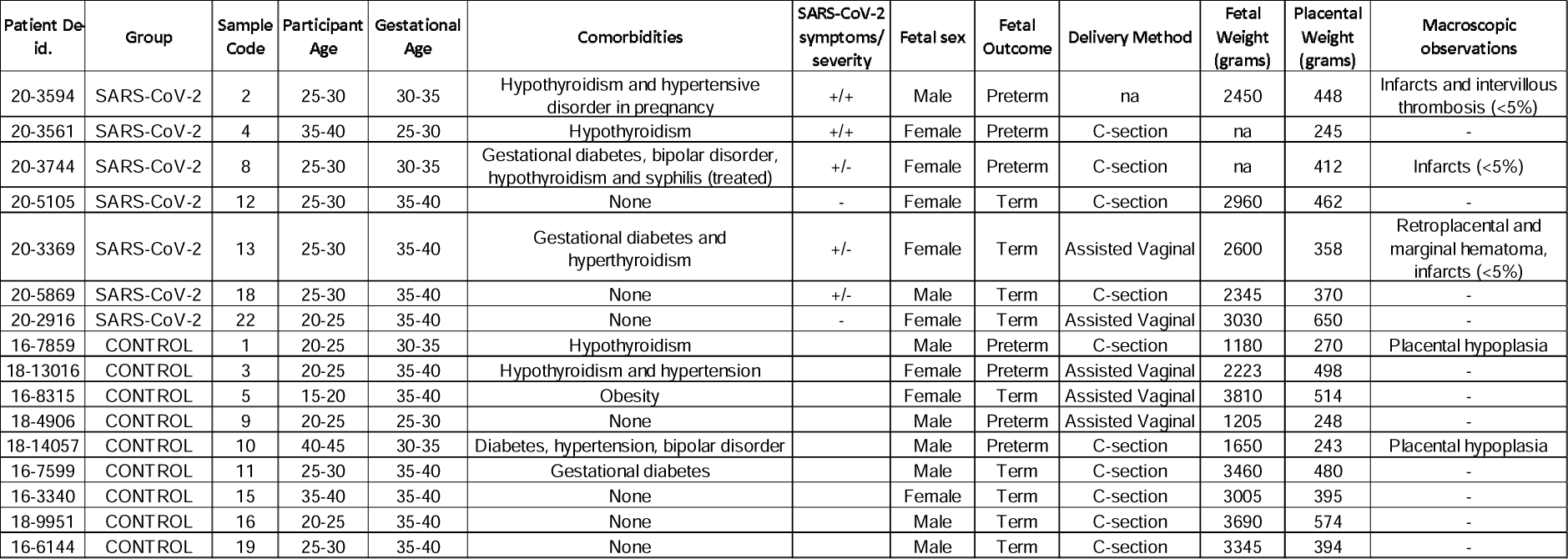
Clinical information of the SARS-CoV-2 and control cohort.

No SARS-CoV-2 viral load was detected in the placental cores from the SARS-CoV-2 infected group through examination by RNAscope of the SARS-CoV-2 spike mRNA (data not shown). With the aid of a trained placental pathologist, an area featuring an anchoring villus, and an area featuring a cluster of terminal villi, were designated as two areas of interest (AOI) within each placental core (Figure 1a). AOIs where immunofluorescently stained with PanCK to identify trophoblast populations, and vimentin to identify stromal populations (i.e., fibroblasts, endothelial cells). Transcriptional expression was collected separately for PanCK positive and separately for vimentin positive cells within each AOI using the Whole Transcriptome Atlas kit (Nanostring; Figure 1a). Subsequent cell deconvolution was performed to assess the purity of each collection (Figure 1b). As expected, transcriptional expression from PanCK positive regions within the AOIs had high proportion of trophoblast populations, compared to vimentin positive regions that had higher proportions of fibroblast and endothelial cells (Figure 1b-f). Due to the overlapping nature of cells, all samples captured a proportion of immune cell types (macrophages, monocytes, Hofbauer cells), except for the PanCK positive regions that displaying a proportion of granulocytes that was absent from the vimentin positive regions within the AOIs (Figure 1b, g-j). SARS-CoV-2 infection did not significantly alter the transcriptional proportion of any cell type assessed when compared to controls (Figure 1b-j). In subsequent analyses, the PanCK positive regions within the AOIs will be referred to as “Trophoblasts” and the vimentin positive regions will be referred to as “Villous Core Stroma” compartments, due to the predominant enriched cell type they represent.

**Figure 1:**
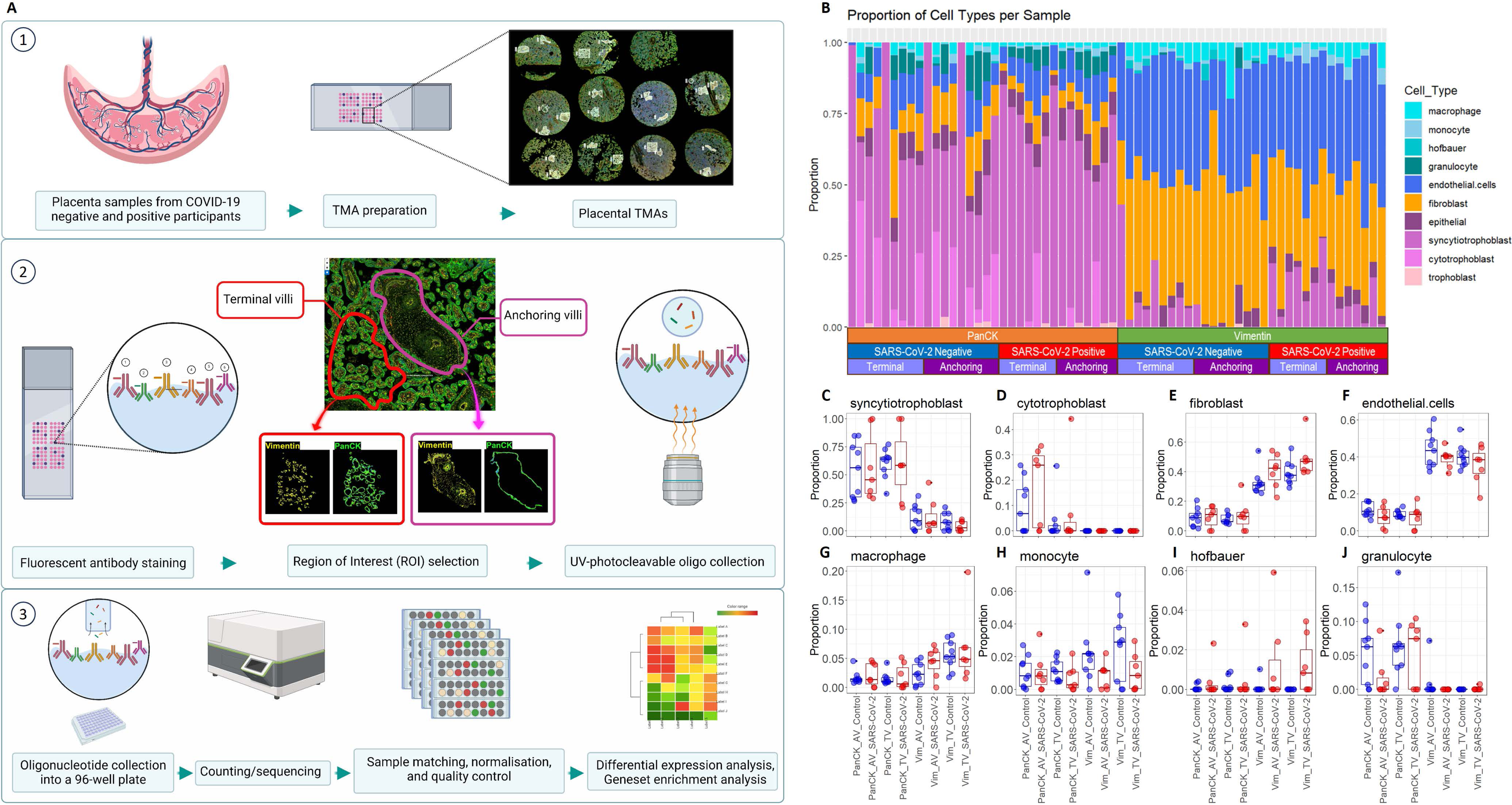
Study design and cell deconvolution. **(a)** 1. Placental cores collected at delivery from the SARS-CoV-2 (n=7) and control (n=9) groups were assembled into tissue microarray slides (TMAs). 2. TMAs were stained with fluorescent markers to differentiate cell types within anchoring (pink outline) and terminal villi (red outline). Barcodes were cleaved and collected from each region of interest by UV light. ***3.*** Cleaved barcodes were sequenced and counted using an Illumina® sequencer in preparation for transcriptomic analysis. Data were normalised before downstream differential expression analysis. **(b)** Transcriptional cell deconvolution map. **(c-j)** Box-plots of indicated cell type proportions from 1b. AV: anchoring villi, TV: terminal villi, SARS-CoV-2 group is n=7 and control group is n=9.

### SARS-CoV-2 infection related pathways enriched in the placenta despite absence of detectable viral particles

Unsupervised clustering of the normalised gene counts by principal component analysis showed that SARS-CoV-2 samples clustered separately to control samples in dimension 1, and further by phenotype in dimension 2, supporting that infection with SARS-CoV-2 markedly alters the transcriptional profiles of the trophoblast and villous core stroma cell populations (Figure 2a-b). Notably, there was very high overlap of genes differentially expressed between the anchoring and the terminal villi; for instance, trophoblasts at the anchoring villi had 1,791 differentially expressed genes versus 493 at the terminal villi, with 405 genes in common between them (82% overlap; Supplementary table 1, Figure 2c). Similarly, the villous core stroma cells at the anchoring villi had 1,139 differentially expressed genes versus 601 at the terminal villi, with 458 genes in common between them (76% overlap; Supplementary table 1, Figure 2c). As expected, there was minimal overlap in differential gene expression between the trophoblasts and villous core stroma compartments, which highlights their distinct cell phenotypes (Figure 2c).

**Figure 2:**
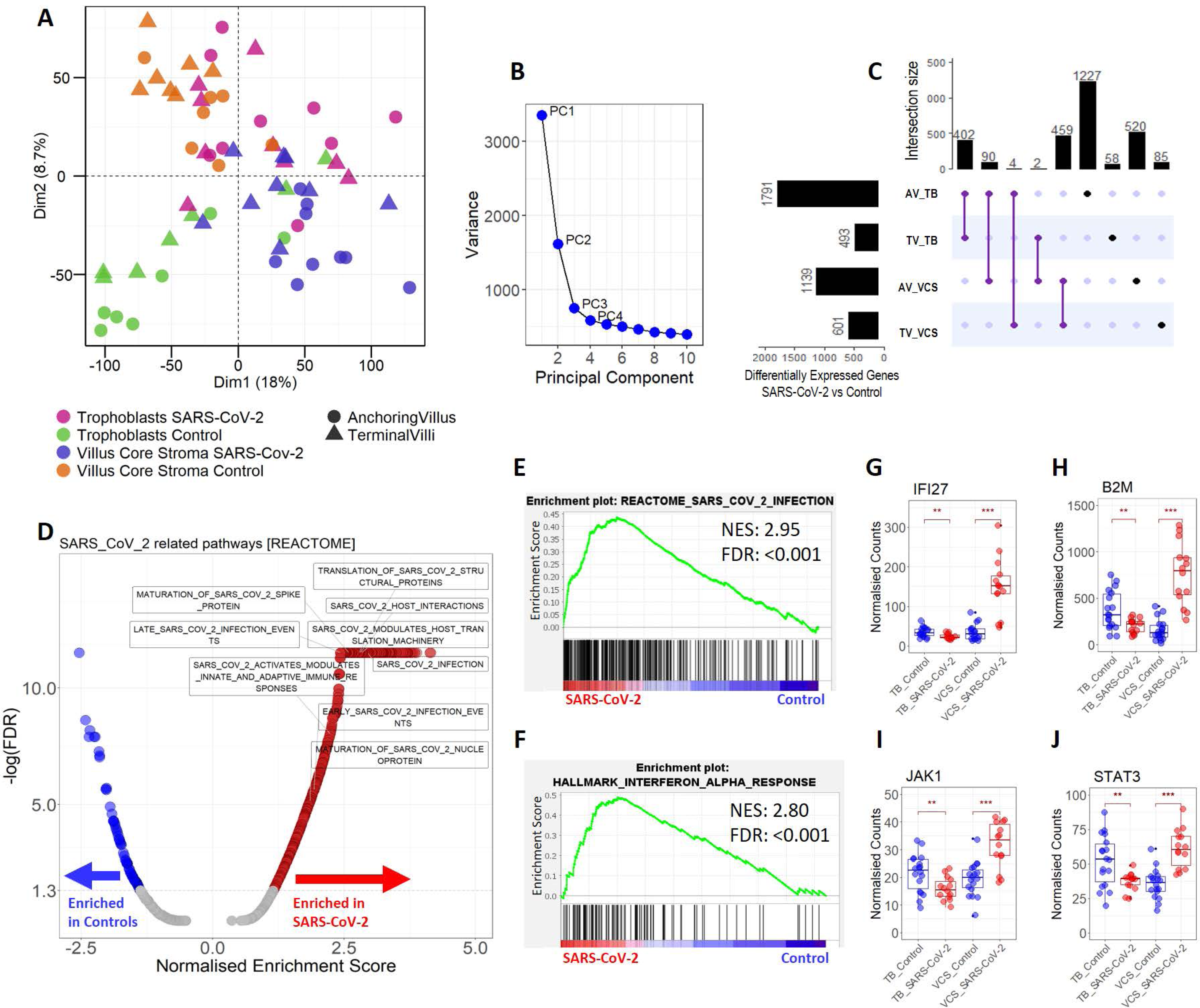
Enrichment of SARS-CoV-2 related pathways. **(a)** Principal component analysis of normalised gene counts from trophoblasts and villous core stromal compartments from SARS-CoV-2 (n=7) and control (n=9) groups at the anchoring or terminal villi (AV; TV). **(b)** Principal component dimensions. **(c)** Upset plot of differential gene expression in trophoblasts and villous core stromal compartments from the AV and TV in response to SARS-CoV-2 infection. The bar charts on the left indicate the total number of differentially expressed genes for the indicated sample group and the bar charts on the top show the gene overlap for the comparisons indicated by the purple lines. Black dots denote differentially expressed genes that are unique for the indicated sample group. Fold change +/- 1.5, *P*-value <= 0.05, FDR < 0.05. (**d)** Enrichment of significant SARS-CoV-2 related pathways from the Reactome database, in the SARS-CoV-2 and control samples. Blue: significantly negatively enriched, red: significantly positively enriched, grey: not significant. The full list of enriched pathways from the Reactome database can be found in Supplementary table 2 and table 3. **(e)** Gene set enrichment analysis (GSEA) plot of the SARS_COV_2_INFECTION pathway from the Reactome database and **(f)** INTERFERON_ALPHA_RESPONSE from the Hallmark database, in SARS-CoV-2 samples versus controls. **(g-j)** Normalised expression counts of IFI27, B2M, JAK1, and STAT3 genes, in trophoblast (TB) or villous core stroma (VCS) compartments from the SARS-CoV-2 (n=7) and control samples (n=9). ** *P*-value < 0.01, *** *P*-value < 0.001.

Despite the SARS-CoV-2 samples showing undetectable SARS-CoV-2 by RNA-scope, transcriptional profiling showed positive enrichment of SARS-CoV-2 related pathways in the SARS-CoV-2 samples, such as “SARS_COV_2_INFECTION”, and “SARS_COV_2_HOST_INTERACTIONS” from the Reactome database, as well as the Interferon Alpha Response pathway from the Hallmark database, which is a first-line immune response pathway that has been associated with SARS-CoV-2 infection (Figure 2d-f)^15^ . These pathways were supported by increased expression of genes that have been associated with SARS-CoV-2, such as the inflammatory marker *IFI16*^16^, disease progression marker *IFI27*^17^, disease prognosticator *B2M*^18^, activation of Janus Kinases (i.e., *JAK1*), and expression of *STAT3*^19^ (Figure 2 g-j). Notably, gene expression for these markers was elevated predominantly in the villous core stroma cell compartment, presumably stemming from the immune subpopulation within the stroma. Indeed, specific analysis of the villus core stroma compartment revealed enrichment of several immune related pathways from the Hallmarks database such as IL6/JAK/STAT3 signalling, IL2/STAT5 signalling, TNF-alpha signalling, inflammatory response, and complement pathways (Figure 3a), supporting that the immune cells within the placental villi are actively responding to SARS-CoV-2 infection.

**Figure 3:**
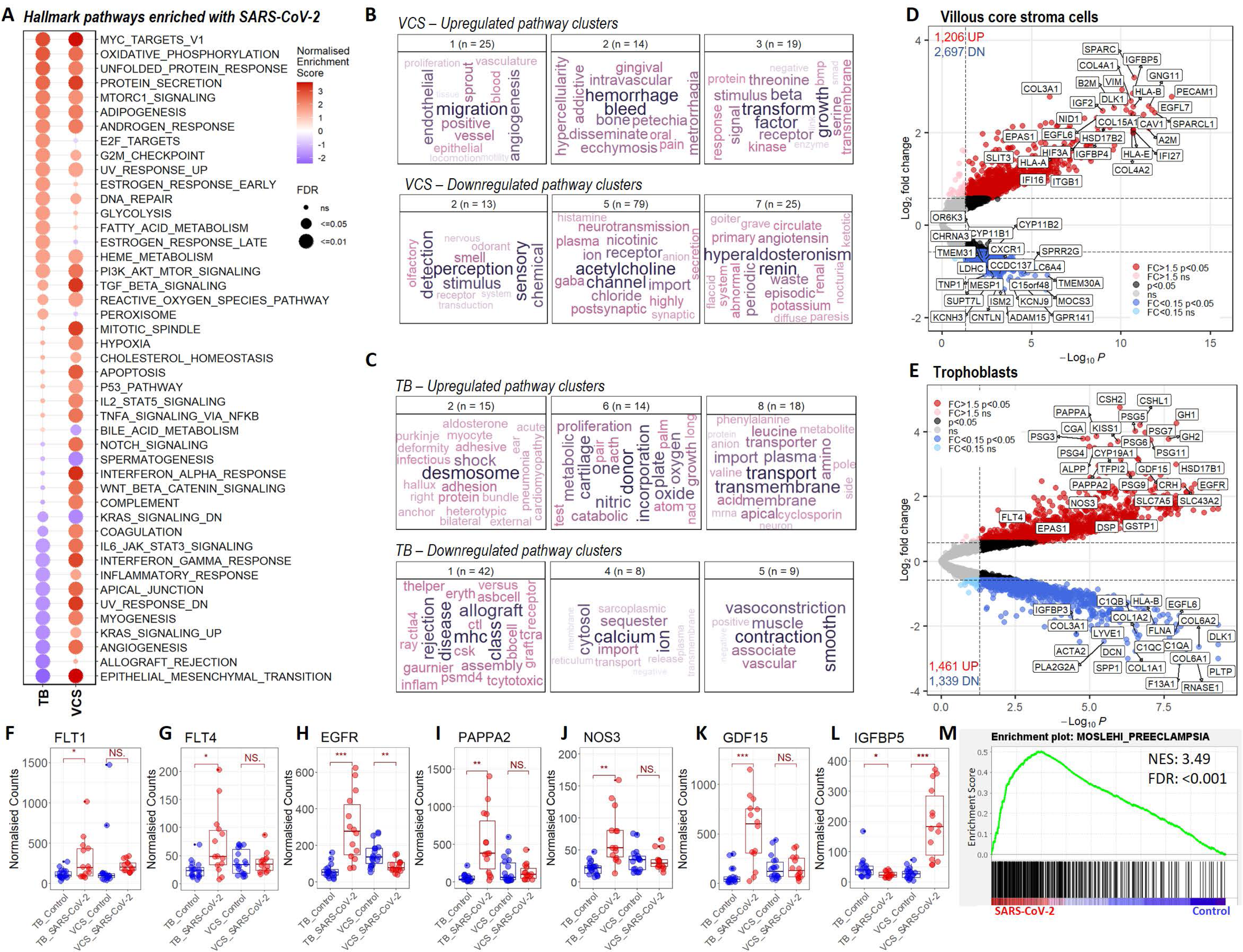
Enrichment of placental dysfunction pathways with SARS-CoV-2. **(a)** Differentially enriched pathways form the Hallmarks database in trophoblasts (TB) and villous core stroma (VCS) compartments in SARS-CoV-2 (n=7) vs control group (n=9). Colour gradient refers to the normalised enrichment score. **(b)** Enriched gene ontology biological processes (GO-BP) pathway clusters from the Molecular Signatures Database (MSigDb) generated using vissE, in the VCS and **(c)** TB. Top row depicts upregulated pathways, bottom row depicts downregulated pathways. **(d)** Volcano plot of gene expression from VCS or **(e)** TB in response to SARS-CoV-2 infection. Fold change (FC) +/- 1.5, *P*-value <= 0.05, FDR < 0.05, the full list of differentially expressed genes can be found in Supplementary table 4. **(f-l)** Normalised expression counts of indicated genes in TB or VCS compartments from the SARS-CoV-2 (n=7) and control samples (n=9). * *P*-value < 0.05, ** *P*-value < 0.01, *** *P*-value < 0.001, NS: not significant. **(m)** Gene set enrichment analysis (GSEA) plot of the preeclampsia signature generated by Moslehi et al, in SARS-CoV-2 samples versus controls.

### SARS-CoV-2 infection enriches hypoxia and placental dysfunction pathways

Pathway enrichment analysis of the genes differentially expressed in response to SARS-CoV-2 infection, revealed pathways related to placental disfunction, in both the trophoblast and villous core stroma compartments. For instance, hypoxia and oxidative phosphorylation pathways were enriched in the villous core stroma (Figure 3a), both of which have been previously linked with placental dysfunction and increased incidence of developing preeclampsia^20^. Hypoxia triggers TGF-β signalling and angiogenesis ^21^, and both TGF-β signalling and angiogenesis related pathways were enriched in the villous core stroma in response to SARS-CoV-2 infection (Figure 3b). Furthermore, pathways related to haemorrhage were upregulated and pathways related to vascular tension, such as hyperaldosteronin/renin pathways, acetylcholine channels, and olfactory receptors, were downregulated in the villous core stroma (Figure 3b). Trophoblast cells exhibited enrichment of pathways related to nitric oxide production (Figure 3c), which is a potent vasodilator^22^. Conversely, pathways related to calcium import^23^ and vasoconstriction were downregulated in trophoblasts (Figure 3c), supporting the notion that the placenta actively reduces vascular tension during SARS-CoV-2 infection. In parallel, trophoblasts showed an increase in cell-cell adherence, communication, and transmembrane amino acid transport, including MTORC1 signalling^24^, suggesting that nutritional uptake to the foetus is enhanced in response to SARS-CoV-2 infection (Figure 3a and 3c). Further, pathways related to allograft rejection and MHC molecules were decreased, suggestive of a defensive mechanism by the trophoblast layer to protect gestation (Figure 3a and 3c).

### Markers associated with preeclampsia are elevated with SARS-CoV-2

Placentae from the SARS-CoV-2 group showed several markers that have been previously associated with hypoxia and placental dysfunction. For instance, the hypoxia and preeclampsia associated markers Fms Related Receptor Tyrosine Kinase 1 (*FLT1) , FLT4*, epidermal growth factor receptor (*EGFR),* and pappalysin-2 (*PAPPA2*) were increased in trophoblasts from the SARS-CoV-2 group (Figure 3e-i)^25–28^. Additionally, markers associated with placental dysfunction and oxidative stress such as Nitric oxide synthase 3 (*NOS3*)^29^, corticotrophin-releasing hormone (*CRH*)^30^, kisspeptin 1 (*KISS1*)^31^, Growth Differentiation Factor 15 (*GDF15*)^32^, and tissue factor pathway inhibitor 2 (*TFPI-2*)^33^, were also elevated in the trophoblasts from the SARS-CoV-2 group (Figure 3e, 3j-k). Transforming growth factor b1 (*TGF*β*-1*), and the PAPP-A2 substrates *IGFBP4/5* (Figure 3d and 3l), were also elevated in the villous core stroma, both markers associated with increased preeclampsia risk^21, 34^.

Given that a number of these pathways and genes are associated with preeclampsia, as well as several recent studies reporting SARS-CoV-2 in predisposing pregnant individuals to preeclampsia^5–10^, we next assessed the enrichment of a preeclampsia-specific gene set generated from published patient cohorts^35^. The gene set was generated by Moslehi et al., where they found 419 genes to be common between four studies examining preeclampsia versus healthy pregnancies^35^. These 419 genes are involved in pathways relevant to preeclampsia, such as oxidative stress, hypoxia, and immune response^35–37^. In our data, this preeclampsia signature was positively enriched in patient samples from the SARS-CoV-2 group (NES 3.49, FDR <0.001; Figure 3m), which aligns with the positive enrichment of hypoxia, immune, and oxidative stress, related pathways we observed in our studies (Figure 3a).

## DISCUSSION

Using digital spatial profiling, we quantified the expression of key markers within distinct cellular compartments of the placenta, providing a detailed picture of the molecular changes occurring in response to SARS-CoV-2 infection. Although our study is limited by its relatively small cohort, cross-sectional nature, and low availability of clinical data, it offers valuable insights into the interplay between trophoblasts and the cells within the villous core stroma in the placenta and how this relationship is influenced by SARS-CoV-2 infection. Close examination of the transcriptional alterations occurring in the placental trophoblast and villous core stroma in response to SARS-CoV-2, revealed a notable number of genes that are enriched in biological pathways previously associated with placental dysfunction.

Trophoblasts from the SARS-CoV-2-infected group had significantly higher levels of *NOS3* compared to the control group (Figure 3j). The upregulation of NOS3 is associated with increased endogenous production of the vasodilator nitric oxide, as a response to altered vascular reactivity, endothelial dysfunction, and hypertension^22, 38^. Interestingly, NOS has been previously found to be highly upregulated to supraphysiological levels in animal models of infection-mediated inflammation during pregnancy, leading researchers to hypothesise that increased NOS may play a role in placental inflammation^39–41^. In response to increased NOS by the trophoblasts, the villous core stromal showed increased expression in biological pathways related to systemic pressure and vasodilation. This included the downregulation of olfactory receptors, acetylcholine channels, and hyperaldosteronin/renin pathways, alongside upregulated hypoxia pathways, suggesting deregulation of the vascular tone and blood pressure due to a hypoxic environment^42^.

Additional transcriptional analysis of trophoblast and villous core stromal compartments from SARS-CoV-2-infected samples identified several transcriptional variations that have been previously associated with preeclampsia (Figure 3). Trophoblasts had higher expression of *EGFR*, a marker that increases with hypoxia and is known to upregulate *FLT1*, where excessive release of soluble FLT1 by syncytiotrophoblasts is a characteristic marker of late onset preeclampsia^43^. There was prominent increase of *PAPPA2* in the trophoblasts, the latter considered to become upregulated in response to hypoxia, and placental pathologies, including preeclampsia^44^. Notably, the PAPPA2 substrates IGFBP4 and IGFBP5 were concurrently upregulated in the villous core stromal compartment, whereby the interaction of PAPPA2 with IGFBP4/5 increases levels of IGF2, which was also increased in the villous core stromal cells in the SARS-CoV-2-infected group (Figure 3d)^34^. Additionally, the villous core stromal compartment had decreased levels of Isthmin-2 (*ISM2*), a placental marker that is downregulated with preeclampsia^45^. *GDF15*, *TFPI-2*, *KISS1,* and *CRH* genes were also upregulated in SARS-CoV-2-infected trophoblasts, all previously associated with placental oxidative stress, hypertension, and preeclampsia^34, 46–48^.

In conclusion, our data suggest that the placenta from pregnancies with SARS-CoV-2 adopts a transcriptional profile aligning with placental dysfunction that has been observed in pregnant participants who develop ‘preeclampsia-like’ syndrome. Using digital spatial profiling, our studies showcased the crosstalk between the trophoblast and villous core stromal cell populations, and how this is enriched with pathways associated with placental dysfunction. Our findings set the foundation for a more comprehensive understanding of placental dysfunction in pregnant individuals with SARS-CoV-2 infection and offer important insights into the potential mechanisms through which SARS-CoV-2 may impact pregnancy outcomes and fetal development.

## METHODS

### Study Design

The SARS-CoV-2 group (n=7) consisted of pregnant, unvaccinated participants, who were symptomatic with COVID-19 in their third trimester (confirmed by RT-qPCR from nasopharyngeal swabs). Placental tissue samples were collected at birth at the Hospital de Clínicas (HC) and Hospital Nossa Senhora das Graças, Brazil, with ethics approval from the National Commission for Research Ethics (CONEP) under approval number 30188020.7.1001.0020^49^. The control group was comprised of archived placentae from nine COVID19-negative people collected during delivery at the Complexo Hospital de Clínicas, Universidade Federal do Paraná, Curitiba, Brazil between 2016 and 2018. To account for maternal co-morbidities, maternal age, and gestational age, the control group was selected to match these clinical features as presented in the SARS-CoV-2 group. Participant cohort and their clinical characteristics are summarised in Table 1. Morphological analysis was performed in all placentas from SARS-CoV-2-infected and control groups using the Amsterdam Placental Workshop Group Consensus Statement^50^. Histological sections were systematically identified and evaluated by two experienced pathologists to obtain samples for tissue microarray (TMA) construction, as described in a previous work^49^. Two TMAs were prepared from the placental samples, following the workflow demonstrated in Figure 1.

### RNAscope

A serial section from the TMAs (4 um) was incubated with RNAscope probes targeting SARS-CoV-2 spike mRNA (nCoV2019, #848568-C3, ACDBio, CA, USA), as per manufacturer’s instructions for automation on Leica Bond RX. DNA was visualised with Syto13 (500 nM, #S7575, ThermoFisher Scientific, MA, USA), and SARS-CoV-2 spike mRNA with TSA Plus CY5 (1:1500, #NEL745001KT, Akoya Biosciences, MA, USA). Fluorescent images were acquired with NanoString GeoMX DSP at 20×.

### Digital spatial profiling with Nanostring GeoMX platform

TMA slides were freshly sectioned (4 um thick serial sections) and prepared according to the Nanostring GeoMX Digital Spatial Profiler (DSP) slide preparation for RNA profiling (NanoString, WA, USA). Slides were hybridised with the NanoString Technologies Whole Transcriptome Atlas (WTA) barcoded probe set (∼18,000 genes), followed by fluorescent staining with Pan-Cytokeratin (PanCK, clone AE-1/AE-3, AF488, Santa Cruz NBP2-33200AF488, [2 µg/mL], CA, USA) to identify trophoblasts, vimentin (VIM, clone E-5, AF594, Santa Cruz sc-373717, [1 µg/mL], CA, USA) to identify endothelial and mesenchymal stromal cells, and SYTO83 to identify nuclei. With the aid of a trained placental pathologist, an area featuring an anchoring villus, and an area featuring a cluster of terminal villi, were designated as two areas of interest (AOI) within each placental core. Oligonucleotides linked to hybridized mRNA targets were cleaved separately for PanCK positive regions within each AOI, and separately for vimentin positive regions. Cleaved oligonucleotides were collected for counting using Illumina i5 and i7 dual indexing as described previously^51, 52^. Paired-end sequencing (2[×[75) was performed using an Illumina NextSeq550 up to 400M total aligned reads. Fastq files were processed using the Nanostring DND system and uploaded to the GeoMX DSP system where raw counts were aligned with their respective AOIs.

### Data normalisation, differential expression analysis, and pathway enrichment analysis

Raw data were normalised to the 134 negative probes in the Human Whole Transcriptome Atlas probe set followed by upper quantile normalisation using the R package RUVseq^53^. Transcriptional data from PanCK positive regions within each AOI were normalised separately to the vimentin positive regions. Differential gene expression analysis between SARS-CoV-2 positive and negative groups was performed separately for PanCK positive regions within each AOI, and separately for vimentin positive regions, using the R package limma^54^. Bayesian adjusted t-statistic method was used where foetal sex and TMA slide number were considered as co-variants. A fold change of +/-1.5 and *P*-value ≤[ 0.05 (adjusted for a false discovery rate of 5%) was considered significant. Pathway enrichment analysis was performed using the Gene Set Enrichment Analysis program (GSEA, v4.3.2, Broad Institute, MA, USA) for biological pathways obtained from the Molecular Signatures Database (MSigDB, Broad Institute, Human v2022.1, MA, USA). The preeclampsia gene-set was obtained from Moslehi et al^35^. GSEA parameters: 1000 permutations, weighted analysis. Gene set enrichment data were further clustered and visualised using the R package vissE with the parameters: computeMsigOverlap (thresh = 0.25), findMsigClusters (alg = cluster_walktrap, minSize =2)^55^.

## AUTHOR CONTRIBUTIONS

**Designing research study**: NS, JR, MA, LdN, AK. **Conducting experiments**: NS, JM, AN, LP, PZR, CMS, ETSS, LdN

**Acquiring data**: JM, AN, LP. **Analyzing data**: NS, IS, NM, AR, LM, GTB, FSFG, VC, AK. **Writing the manuscript**: all authors

## ACKNOWLEGMENTS

This study was funded by the Queensland University of Technology ECR funds for AK, JR, MA, NS. The following authors are supported by fellowships from the NHMRC (AK – 1157741, GB - 2008542 and 1135898), US DOD (NS – PC190533). The authors thank Fred Hutch pathology (Miki Haraguchi & Stephanie Weaver) for histology assistance.

## DATA AVAILABILITY

The data generated in this study are available in the Gene Expression Omnibus (GEO) under GSE223612.

